# Hair test reveals plasticity of human chronotype

**DOI:** 10.1101/2025.03.07.641864

**Authors:** Bert Maier, Luísa K. Pilz, Selin Özcakir, Ali Rahjouei, Ashraf N Abdo, Jan de Zeeuw, Dieter Kunz, Achim Kramer

## Abstract

Circadian clocks govern daily physiological and behavioral processes and are crucial for health, yet disruptions can lead to various diseases. Chronotype, the state of circadian timing, varies between individuals and is reflected in behaviors such as sleep-wake patterns, cognitive performance, and physical activity. This interindividual variability is influenced by both genetic factors and environmental cues, but the relative contributions of each remain unclear, particularly in terms of plasticity - how much chronotype can shift in response to lifestyle and environmental factors. The gold standard for chronotype assessment, dim-light melatonin onset (DLMO), is invasive and impractical for large-scale studies, while blood-based molecular biomarker tests, which estimate internal time, show promise but are limited by practicality. Here, we introduce HairTime, a novel assay that estimates chronotype from a single hair sample collected at one point during the day. HairTime was developed and evaluated in two studies: a training study and a validation study, where it demonstrated a strong ability to predict chronotype, with DLMO as the comparison. This non-invasive method is suitable for large-scale, longitudinal studies and clinical practice. We assessed HairTime using over 4,000 samples, observing a normal distribution of chronotype across the population, with its estimation associating with age, sex, and notably, work schedules. The association with work schedules reveals the plasticity of chronotype, as workdays circadian timing earlier, highlighting that societal factors can influence and modify an individual’s internal rhythm. Additionally, we explore the concept of circadian amplitude, finding that lower amplitude rhythms in hair follicle cells are linked to reduced chronotype prediction accuracy. Our results highlight that both intrinsic circadian mechanisms and external factors, such as lifestyle and work schedules, shape chronotype. HairTime offers an innovative tool for understanding circadian rhythms, facilitating personalized chronotherapy to improve health outcomes by aligning treatments with an individual’s biological rhythms.

## Introduction

Human daily behavior is regulated by an endogenous timekeeping system known as the circadian clock. This genetic program is present in nearly all human cells, where individual cellular oscillators form networks that collectively constitute the circadian system. As a result, daily rhythms emerge at the cellular, organ, and whole-organism levels. The circadian system orchestrates physiological and behavioral processes with respect to time of day and uses environmental cyclic cues, known as zeitgebers, to synchronize the body’s internal time with the external environment—a process called entrainment. In mammals, the principal pacemaker, or central clock, is located in the suprachiasmatic nucleus (SCN) in the hypothalamus. The SCN is primarily entrained by the light-dark cycle and transmits time-of-day information to oscillators in peripheral tissues (Foster, 2020; Takahashi, 2017). Through this entrainment process, circadian rhythms establish specific phase relationships with external zeitgeber cycles, known as the phases of entrainment. In humans, there is considerable individual variation in these phases of entrainment, leading to inter-individual differences in the timing of clock-controlled functions such as hormone secretion, cognitive performance, and physical activity. The sleep-wake cycle is one of the most prominent physiological processes governed by the circadian system. These individual differences in entrainment phase—referred to as *chronotypes*—are most evident in sleep patterns, where some individuals have naturally earlier sleep times (“larks”) while others have later sleep times (“owls”). Traditionally, chronotype is assessed using questionnaires, such as the Munich ChronoType Questionnaire (MCTQ) and the Morningness-Eveningness Questionnaire (MEQ) (Horne & Ostberg, 1976; Roenneberg et al., 2007). For a more objective assessment, various methods have been proposed, but the current gold standard is the measurement of dim-light melatonin onset (DLMO)—the point at which the pineal gland begins melatonin production in the absence of light. Since melatonin secretion is predominantly regulated by the SCN, DLMO serves as a reliable marker for the onset of the biological night (Klerman et al., 2022).

It is becoming increasingly evident that synchronization between endogenous circadian rhythms and external environmental cycles is essential for health and well-being. However, modern lifestyles including shift work, travel across time zones or irregular mealtimes challenge our internal clock, often leading to circadian disruption (De Assis & Kramer, 2024; Finger & Kramer, 2021). Today, it is widely accepted that such disruptions are linked to a wide range of common diseases, including sleep disorders, psychiatric and neurodegenerative conditions, metabolic and cardiovascular diseases, immune dysfunctions, and even cancer. Despite recent advances, the crucial interplay between the circadian clock and health remains largely underutilized in medical education and practice. For instance, it is well established that disease symptoms (Scheer et al., 2021) and medical emergencies such as asthma attacks and heart attacks (Cohen et al., 1997; Muller et al., 1985) follow distinct daily patterns. Additionally, many genes encoding druggable proteins exhibit cyclic transcription in at least one tissue in primates (Mure et al., 2018). Yet, circadian timing is rarely considered in drug development, even though studies suggest that aligning treatments with biological rhythms can enhance effectiveness and reduce side effects (Abusamak et al., 2025; Pigazzani et al., 2024). A new field, circadian medicine (or chronomedicine), is now emerging to bridge this gap. It aims to uncover the mechanisms linking the circadian clock to health and disease and apply this knowledge to improve diagnosis, treatment, and prevention (Kramer et al., 2022).

Circadian precision medicine aims to optimize treatment by aligning medical interventions with an individual’s biological rhythms. As mentioned earlier, people vary in their chronotypes, making the assessment of internal time crucial for maximizing the benefits of circadian medicine. The current gold standard for estimating chronotype, DLMO, is labor-intensive and requires multiple saliva or blood samples collected in the evening ideally under highly controlled conditions (Murray et al., 2024). These constraints make DLMO assessment impractical for large-scale studies and routine clinical applications. To circumvent these challenges, self-reported questionnaires such as the MCTQ and the MEQ are commonly used. The MCTQ estimates chronotype based on reported sleep-wake times, while the MEQ assesses individual preferences for daily activities and self-perceived states (e.g., alertness, tiredness) at different times of day. However, both rely on subjective self-assessment, which introduces potential biases. Several factors beyond the circadian clock may influence questionnaire-based chronotype estimates. For instance, individuals with extreme chronotypes who struggle with rigid societal schedules may report their preferred timing on the MEQ reflecting their wish to have more moderate chronotypes. Similarly, MCTQ-based estimates can be affected by sleep homeostasis factors, such as sleep debt, despite computational adjustments designed to minimize this influence.

Recently, several research groups have developed more feasible objective molecular biomarker tests that require only a single or a few blood or tissue samples. Gene expression in peripheral tissues can serve as a marker of the human circadian clock’s state and function (Braun et al., 2018; Wu et al., 2018), as the molecular clock machinery driving endogenous rhythms is present in almost all tissues (Mure et al., 2018; Takahashi, 2017) and is synchronized by the SCN. These advancements may enable objective chronotype determination in larger cohorts (Dijk & Duffy, 2020; Münch & Kramer, 2019). We developed BodyTime, an assay that estimates internal time from a single blood sample using a small set of transcript biomarkers (Wittenbrink et al., 2018). Compared to DLMO, BodyTime demonstrated a strong performance while offering several advantages: (i) Simpler integration into study protocols and clinical settings; (ii) Lower cost and reduced logistical constraints (e.g., no need for controlled conditions during sample collection); (iii) Objective measurement, avoiding the subjectivity of self-reported questionnaires. However, BodyTime has technical challenges that limit its scalability, particularly in large-scale studies and field research. The requirement to keep blood cells cold and isolate monocytes makes its widespread application difficult. To address these limitations, hair follicle cells were identified as a promising, less invasive alternative biospecimen. In these cells, clock gene expression follows a daily rhythm, with its phase aligning with behavioral rhythms (Akashi et al., 2010). Additionally, because hair follicle cells remain attached to plucked hairs, no cell separation is required, and the RNA yield from a few plucked hairs is sufficient for standard gene expression analysis (Kim et al., 2006). A hair-based assay would facilitate internal time assessment in large-scale and field studies, enable longer follow-up periods for study participants and outpatients, and allow for more frequent sampling in longitudinal studies.

Chronotype, or the phase of entrainment, is a state shaped by the interaction between the endogenous circadian clock and the environmental zeitgeber cycle. While genetic differences explain why individuals synchronize differently to the same light-dark cycle, chronotype also depends on zeitgeber strength (Granada et al., 2013). For instance, when individuals transition from urban environments to natural settings—where daylight exposure is high during the day and minimal at night—their chronotype shifts significantly earlier (Wright et al., 2013). However, the extent to which chronotype fluctuates in response to environmental factors such as seasonal changes, weather conditions, and daily behavioral variations remains largely unknown. A key limitation in addressing this question is the lack of practical methods for longitudinal chronotype assessment. Recent findings suggest that shifts in dim-light melatonin onset (DLMO) can occur over even shorter timescales, with measurable delays on work-free days compared to workdays (Zerbini et al., 2021). In modern lifestyles, characterized by irregular light exposure and inconsistent mealtimes, understanding chronotype stability is crucial—not only for circadian research but also for personalized circadian medicine and preventive strategies. Helping individuals live in alignment with their internal clocks could reduce the risk of circadian disruption, which is increasingly linked to common chronic diseases (De Assis & Kramer, 2024).

In this study, we developed an assay (HairTime) to estimate chronotype from hair samples collected at a single time point and validated it against the gold-standard marker for circadian phase of entrainment, dim-light melatonin onset (DLMO). Compared to blood sampling, hair collection is minimally invasive and does not require a healthcare professional—participants can collect their own samples and mail them in. This makes the assay easy to implement, cost-effective, and well-suited for large-scale studies and longitudinal assessments. We demonstrate HairTime’s applicability by analyzing chronotype estimates from hair follicle samples in a large cohort of over 4,000 adults. Our findings show that hair-derived chronotype is normally distributed in the German population and varies with age and sex. Furthermore, we investigated the plasticity of human chronotype by exploring the relationship between hair-derived chronotype and work schedules. Our results suggest that chronotype is influenced by the number of workdays, shedding light on how societal demands shape the circadian system and highlighting the impact of modern work-life patterns on our natural rhythms.

## Results

To identify genes that are expressed in human hair roots in a time-of-day dependent manner, we first performed a pilot study with three healthy volunteers. The objective was to identify rhythmic candidate genes in a small cohort, which could then be used in a larger study to build a training dataset for chronotype predictions from a single sample, without the necessity of analyzing all genes in the second study. The test subjects pulled out some of their own hair from the scalp at regular four-hour intervals over a period of 36 hours. The RNA from the hair root cells was extracted, and the gene expression was analyzed by RNA sequencing. A total of 90 genes that exhibited consistent rhythmic activity were selected (**Supplementary Table 1**).

To generate a training dataset, we conducted the *HairTime study*, in which we analyzed the gene expression of the 90 selected genes in conjunction with six housekeeping genes in 13 healthy male and female volunteers using a 96-plex NanoString panel (**Supplementary Table 2 and 5**). The volunteers were instructed to sample their hair roots at regular three-hour intervals over a 33-hour period (**Figure 1A**). To timestamp these data with respect to the individual’s internal time, we also determined the dim-light melatonin onset (DLMO) from saliva samples of each participant on the second day of the study. This enabled us to describe the gene expression of the 90 candidate genes as a function of internal time (**Figure 1B, Supplementary Table 3**).

**Figure 1.**
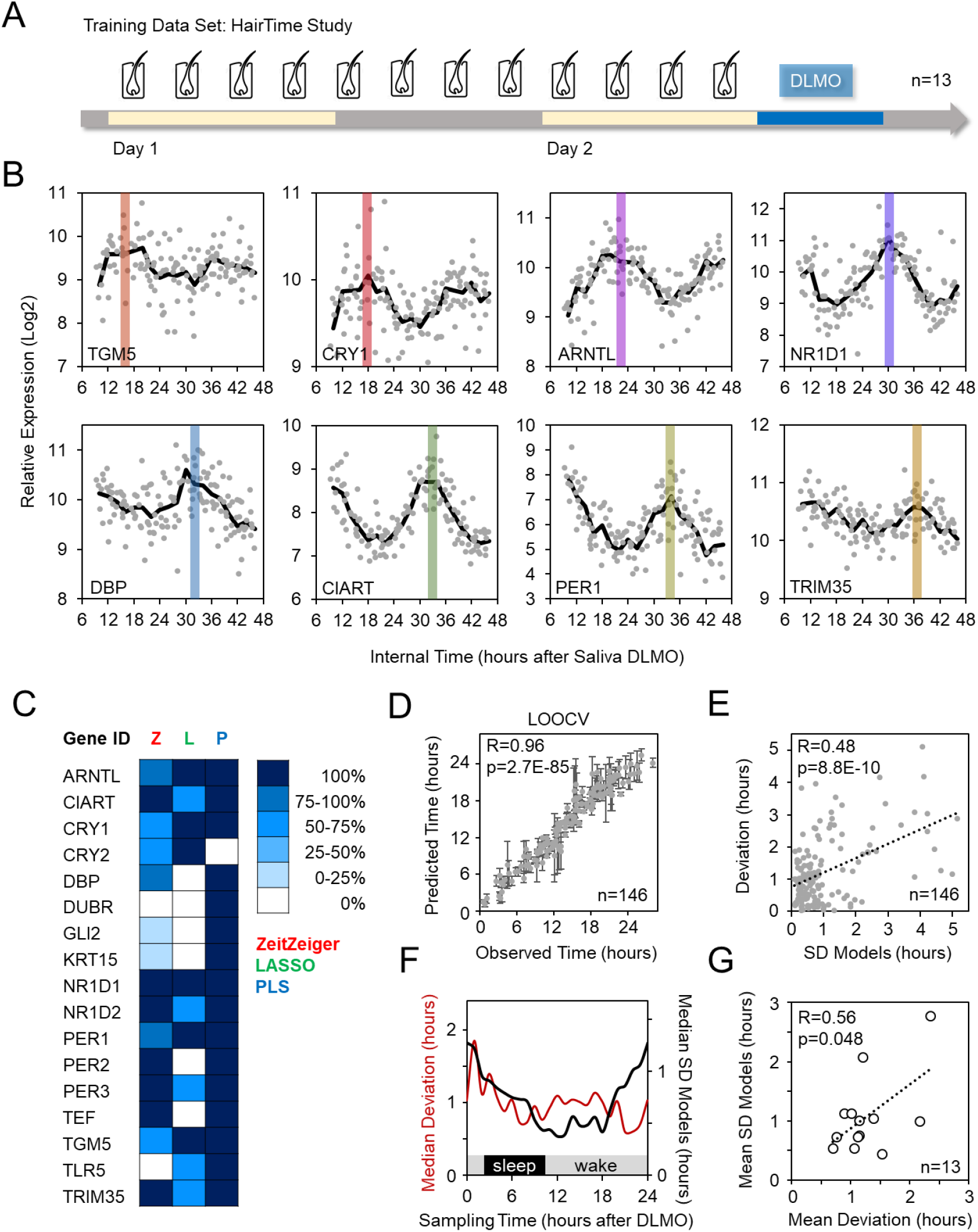
Identification of transcriptional biomarkers from hair roots for prediction of human chronotype. (**A**) HairTime study setup. (**B**) Composite expression rhythms of the indicated genes from the hair root samples of all 13 participants in the HairTime study. The colored bars indicate the peak expressions of the genes. (**C**) Selected features (genes) and their relative frequency of occurrence in the different parameter versions of the ZeitZeiger, LASSO, and PLS models. (**D**) Pearson’s correlation between the observed internal time and the predicted internal time (mean of the three model predictions) from a leave-one-out cross-validation (LOOCV) approach. Error bars represent the standard deviation (SD) of the model predictions. (**E**) Pearson’s correlation between the degree of model disagreement (models SD) and the deviation of the hair prediction from the saliva DLMO value. (**F**) Prediction accuracy and model agreement as a function of sampling time (given in hourly bins). Prediction accuracy is given as median deviation from saliva DLMO value for all samples per bin. Model disagreement is given as median of the SD of model predictions per bin. (**G**) Pearson’s correlation between mean deviation of the predictions from saliva DLMO value and the disagreement of the models (given as mean SD of models) per study participant.

To identify the most effective genes for predicting internal time (DLMO) from a single sample, we applied three machine learning methods—ZeitZeiger, LASSO, and partial least squares (PLS)— combined with a leave-one-out cross-validation approach (see Methods for details). For each method, we selected two models that achieved optimal prediction accuracy while minimizing the number of required genes. To ensure a balanced estimate and mitigate potential biases, chronotype predictions were averaged across the selected ZeitZeiger, LASSO, and partial least squares models. In total, 17 genes were identified for chronotype prediction (**Figure 1C, Supplementary Table 4**).

While the prediction accuracy was remarkably high (median absolute deviation [MdAE] from saliva DLMO of approximately one hour, **Figure 1D**), nearly 20% of samples in the training dataset exhibited a prediction deviation of more than two hours. Interestingly, in these samples, the three prediction models frequently did not agree. Indeed, the standard deviation of the three predictions demonstrated a highly significant correlation with the discrepancy between the prediction and the saliva DLMO (**Figure 1E**). Consequently, the standard deviation of the model predictions should enable the estimation of the accuracy of a prediction for future, hitherto unknown samples.

What factors contributed to the discrepancy in predictive accuracy across samples? To explore potential causes, we conducted a detailed analysis to assess whether prediction accuracy varied by time of day and whether there were notable interindividual differences among participants. Indeed, the prediction models exhibited considerable inconsistencies when using samples collected between approximately four hours before and eight hours after DLMO (i.e., during the evening and nighttime), and were particularly inaccurate when preformed on nighttime samples (**Figure 1F**). Moreover, we observed substantial interindividual differences in mean prediction accuracy, which correlated significantly with the mean standard deviation of the prediction models across all samples from a given individual (**Figure 1G**). This provides further support for the hypothesis that discrepancies between prediction models may serve as an indicator of accuracy.

Why does prediction accuracy vary between individuals? We hypothesized that this could be linked to the amplitude of their gene expression rhythms This assumption is plausible, as lower amplitudes in noisy expression data are expected to increase susceptibility to errors in prediction models and lead to greater inconsistencies between models using different approaches. To test this hypothesis, we analyzed the amplitude of gene expression rhythms for each study participant. As a result of the significant positive correlation between the amplitudes of numerous gene expression rhythms (**Figure 2A,B**), an amplitude score was developed as an indicator of the amplitude of the circadian clock machinery in hair root cells. This score was derived from the five genes that exhibited the highest amplitude and strongest cross-correlation (*CIART, NR1D1, NR1D2, PER3, TEF*). Indeed, individuals with lower amplitude scores demonstrated a significantly lower degree of agreement between the predictions of the three models (**Figure 2C**). This finding suggests that the accuracy of chronotype prediction from a single biospecimen is dependent on the strength of a person’s circadian system.

**Figure 2.**
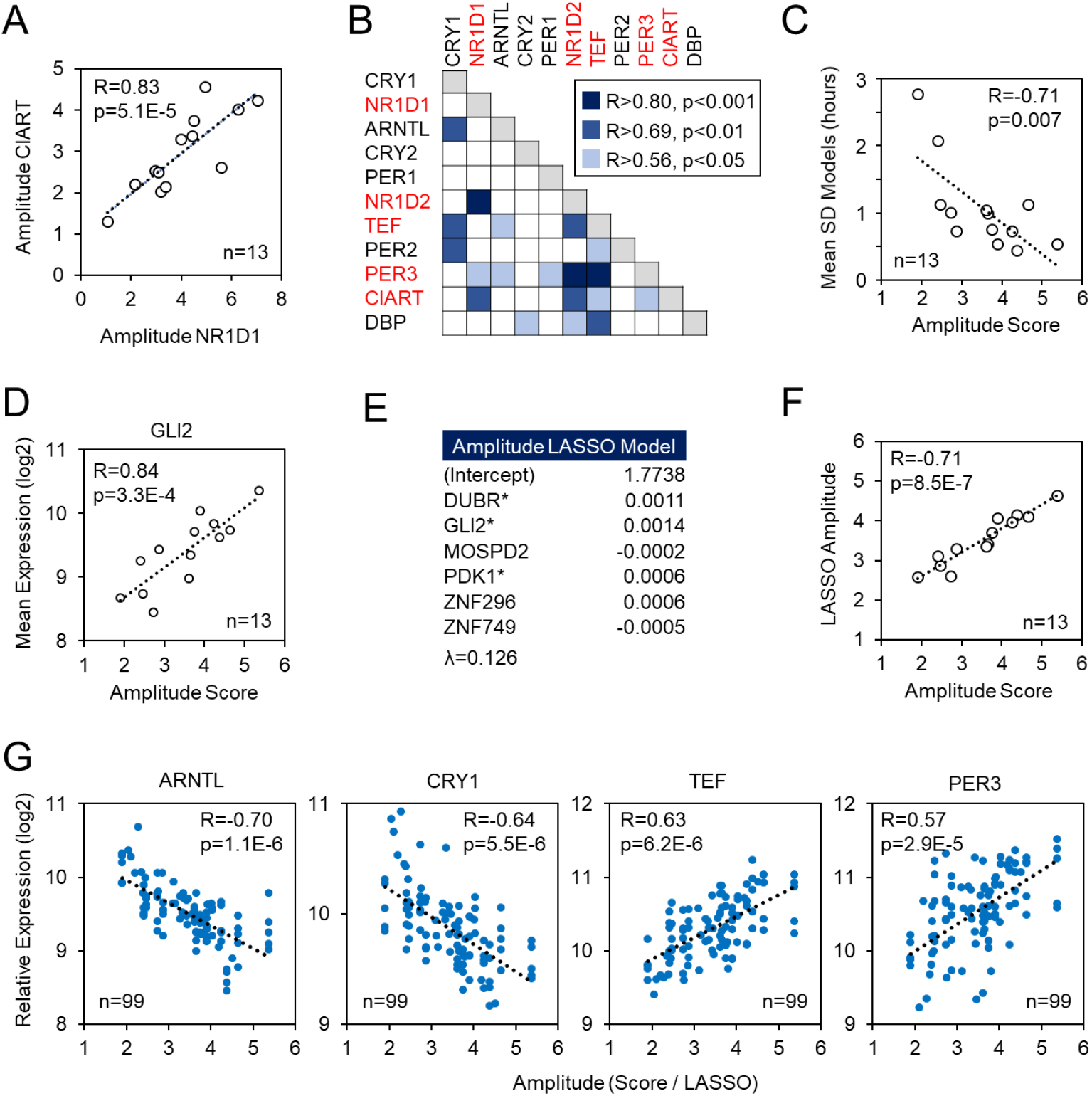
Prediction of circadian amplitude using transcriptional biomarkers. (**A**) Pearson’s correlation between rhythmic expression amplitudes of *NR1D1* and *CIART* in the 13 HairTime study participants. (**B**) Pairwise Pearson’s correlation between the amplitudes of the expression rhythms of the indicated genes (red: strongest cross-correlation). (**C**) Pearson’s correlation between the amplitude score, calculated as the geometric mean of the expression rhythm amplitudes of the genes indicated in red in (B), and the mean model disagreement for each HairTime study participant. (**D**) Pearson’s correlation between the amplitude score and the mean expression of the gene GLI2. (**E**) LASSO model coefficients for predicting circadian amplitude derived from 46 non-rhythmic candidate genes. Parameter λ was adjusted to limit the number of predictive biomarkers to a maximum of six. Expression levels of genes marked with asterisks show a significant correlation with the amplitude score (as shown in (D)). (**F**) Pearson’s correlation between amplitude scores and predicted amplitudes from LASSO model shown in (E)). (**G**) Pearson’s correlation between gene expression level and amplitude. Shown are morning to noon samples from HairTime study participants along with samples from HairVali study participants (see Figure 3A). Amplitudes are either calculated amplitude scores (for HairTime) or LASSO predicted values (for HairVali).

It is frequently observed that predictors demonstrate superior performance on data sets on which they were constructed than on independent samples. Consequently, it is critical to validate them externally prior to implementation for clinical or research purposes. Thus, to validate the final predictors, an independent study (*HairVali study*) with 35 healthy male and female volunteers was conducted (**Supplementary Table 2**). The saliva DLMO and the expression of the 17 prediction genes from a single hair root sample taken between 9 and 17 hours after DLMO (i.e., between morning and early afternoon) were determined (**Figure 3A**). In this independent data set, the prediction accuracy of DLMO from hair root gene expression using the three models was again remarkably high, with a combined MdAE of 0.99 h and more than 77% of all samples exhibiting a prediction deviation of less than 2 hours (**Table 1, Figure 3B**). However, some samples exhibited markedly inferior prediction accuracy, prompting us to investigate potential causes. As with the training dataset, we observed a significant correlation between the discrepancy between model predictions and actual DLMO and the deviation among the predictions themselves. This suggests that the poorly predicted samples may be associated with a low amplitude of the individual’s circadian system (**Figure 3C**).

**Figure 3.**
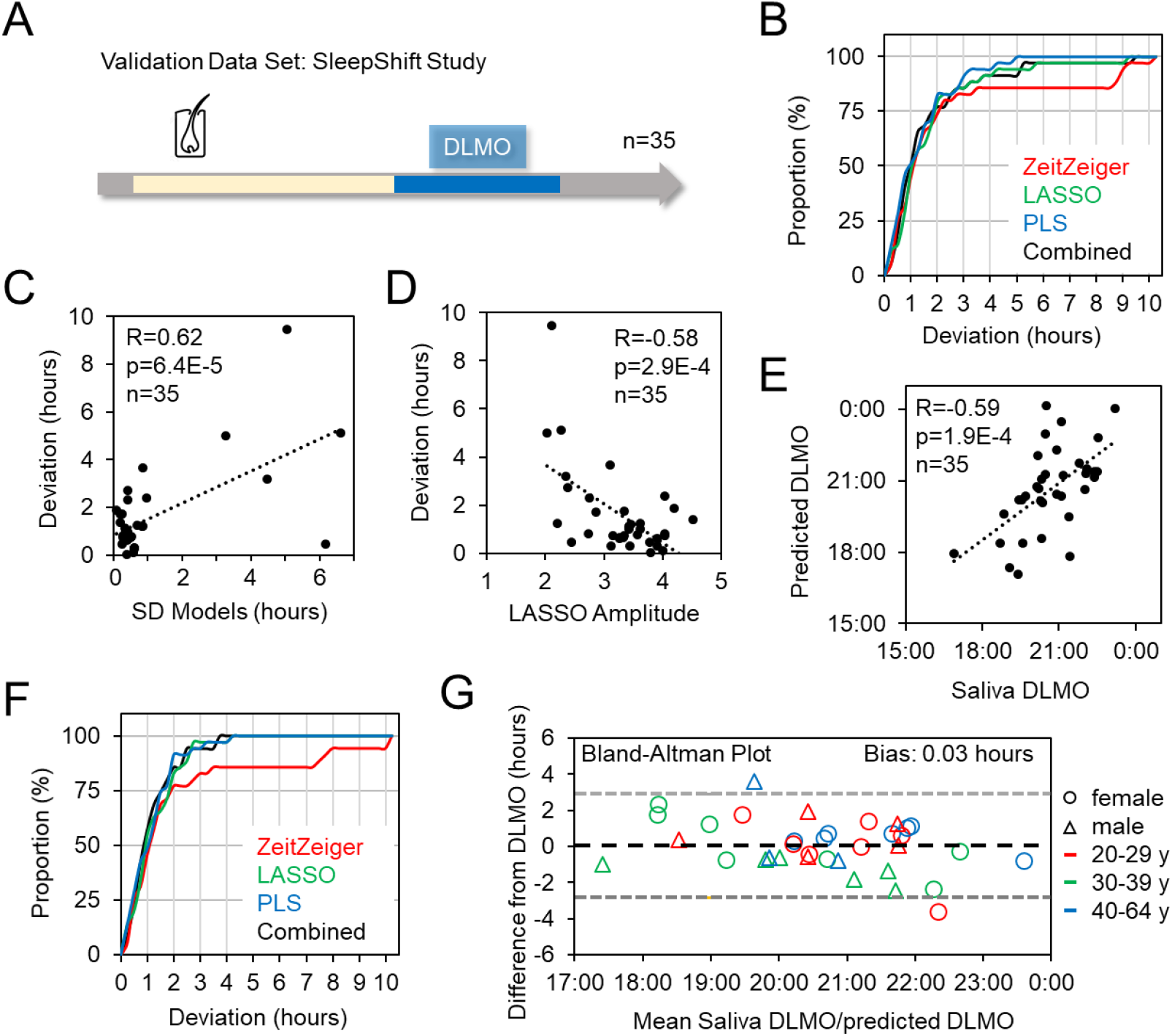
Validation of biomarkers for prediction of human chronotype in an independent study. (**A**) HairVali study setup. (**B**) Cumulative frequency distributions of the absolute prediction deviation from saliva DLMO values for three different models and their mean (combined) when applied to the HairVali study data set. (**C**) Pearson’s correlation between model disagreement and deviation from saliva DLMO values for samples of the HairVali study. (**D**) Pearson’s correlation between the LASSO-predicted amplitude of the circadian system of participants in the HairVali study and the deviation of the predicted DLMO from the saliva DLMO value. (**E**) Pearson’s correlation between saliva DLMO and DLMO predicted from hair root biomarkers for HairVali study samples. Gene expression values of six low-amplitude samples (LASSO amplitude < 2.4) were adjusted as described in the main text and in the Materials and Methods section. (**F**) Cumulative frequency distributions of the absolute prediction deviation from saliva DLMO values for three different models and their mean (combined) when applied to the HairVali study data set after adjustment of low-amplitude samples. (**G**) Bland-Altman analysis of the bias between saliva DLMO and DLMO predicted from hair root biomarkers (combined). Black dashed line corresponds to mean difference between methods; grey dashed lines correspond to the limits of agreement (mean difference ± 1.96 SD).

**Table 1:**
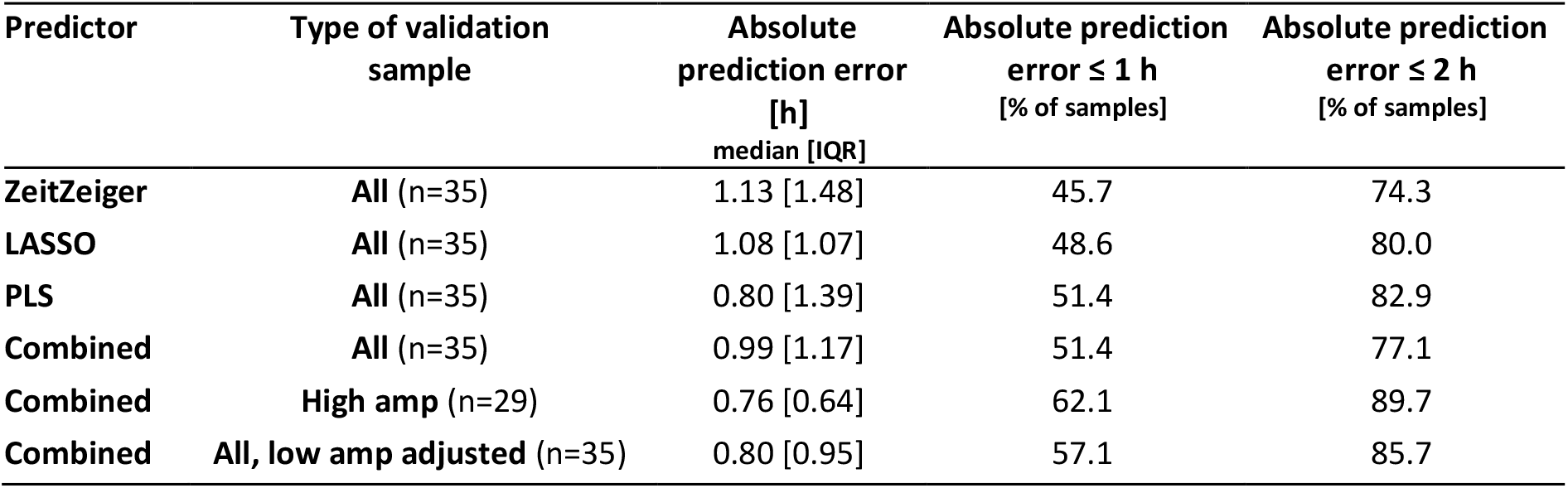
External validation of the HairTime predictors in the independent HairVali study.

| Predictor | Type of validation sample | Absolute prediction error [h]<br>median [IQR] | Absolute prediction error ≤ 1 h [% of samples] | Absolute prediction error ≤ 2 h [% of samples] |
| --- | --- | --- | --- | --- |
| **ZeitZeiger** | **All** (n=35) | 1.13 [1.48] | 45.7 | 74.3 |
| **LASSO** | **All** (n=35) | 1.08 [1.07] | 48.6 | 80.0 |
| **PLS** | **All** (n=35) | 0.80 [1.39] | 51.4 | 82.9 |
| **Combined** | **All** (n=35) | 0.99 [1.17] | 51.4 | 77.1 |
| **Combined** | **High amp** (n=29) | 0.76 [0.64] | 62.1 | 89.7 |
| **Combined** | **All, low amp adjusted** (n=35) | 0.80 [0.95] | 57.1 | 85.7 |

To study circadian amplitude, time series data from individuals are required, as this parameter cannot yet be estimated from a single sample. To address this limitation, we systematically searched our training data for genes whose absolute expression levels could predict amplitude, as previously described in mouse models (Littleton et al., 2020). Using LASSO regression, we modeled amplitude scores from HairTime participants as the independent variable and gene expression levels from morning to noon as the dependent variables (**Figure 2D,E**). To minimize the influence of temporal fluctuations, our analysis focused on 46 of the 90 trainig data set genes with the least variability in expression (not shown). A model using six genes (*DUBR, GLI2, MOSPD2, PDK1, ZNF296*, and *ZNF749*) was identified for amplitude prediction. The predicted amplitudes based on these genes showed a significant correlation with measured amplitude values, providing confidence in their applicability for predicting amplitudes in the 35 HairVali sample (**Figure 2F**).

Utilizing this amplitude prediction model, we calculated amplitude scores for the 35 HairVali participants, which ranged from 2.0 to 4.5 (mean 3.3, 95% CI [3.05; 3.50]). This is analogous to the observations made for the 13 HairTime participants, for whom the range was 1.9 to 5.4 (mean 3.5, 95% CI [2.92; 4.41]). As anticipated, the prediction accuracy of the 35 HairVali samples exhibited a significant correlation with the amplitude scores derived from the model. The prediction error was higher for samples from participants with low amplitudes than for samples from participants with normal or high amplitudes (**Figure 3D**). Subsequently, the accuracy of our phase prediction was analyzed without the samples from subjects with low amplitude (6 out of 35). The resulting MdAE was 0.76 hours, with 90% of all samples exhibiting less than a 2-hour deviation and 62% of all samples deviating by less than an hour from the measured DLMO (**Table 1**).

Although amplitude prediction can be used to identify samples whose phase prediction is likely to be less accurate, our next objective was to obtain more accurate phase prediction for low-amplitude samples. In our previous study with septic shock patients (Lachmann et al., 2021), we observed that the amplitudes of the clock gene expression rhythms were markedly low, and that the absolute mean expression levels of these genes were frequently significantly altered. We thus sought to determine whether such an association could also be identified in our healthy subjects. Indeed, a significant positive correlation was identified between amplitude and mean expression level for nine of the 17 phase prediction genes (*CIART, CRY2, DBP, NR1D2, PER1, PER2, PER3, TEF*, and *TRIM35*) (**Figure 2G**). To account for this relationship, the gene expression values of the six low-amplitude samples from the HairVali study were adjusted using the regression parameters derived from the linear correlations. This adjustment ensured that the expression values corresponded to those of a sample with a mean amplitude score of 3.5 (see Materials and Methods for details). The application of these adjusted expression values to the phase prediction of the six previously described samples resulted in a notable improvement in accuracy and a highly significant correlation between the DLMO predicted using a single hair sample and the DLMO measured in saliva (**Figure 3E**). Consequently, the overall prediction accuracy of the 35 samples (including the 6 adjusted) is MdAE = 0.80 hours, with 86% of all samples having less than a two-hour deviation and 57% of all samples having less than an hour’s deviation from the measured DLMO (**Table 1, Figure 3F**). In addition, Bland-Altman analysis reveals no systematic bias between the hair-based prediction and DLMO (mean difference 0.03 hours, 95% CI [-2.84; 2.90]) (**Figure 3G**).

The combined results demonstrate that our novel objective chronotype test is capable of accurately predicting the internal phase from a single hair sample. Furthermore, the amplitude score and the application of three distinct models provide a means of estimating the accuracy of the prediction. In instances where the prediction is deemed to be inaccurate, the method allows for the correction of low-amplitude samples, thereby enhancing the precision of the phase prediction.

To characterize chronotype predicted from hair in a larger cohort and assess whether the values align known distributions and epidemiological patterns in the field, we analysed data from the BodyClock dataset. Hair samples were collected from 4,351 individuals between December 30, 2021, and June 21, 2024, with most participants located in Germany at the time of sampling. Socio-demographic and occupational data were available for a subset of the sample (n = 4,044). We excluded individuals younger than 18 or older than 70, as these age groups were underrepresented and less likely to yield reliable estimates, resulting in a final sample size of n = 3,949 for analyses examining associations between age, sex, and work status with hair-based DLMO predictions. See **Supplementary Table 6** for characteristics of each sample.

The distribution of hair follicle–predicted chronotype (pDLMO) was visually assessed and found to be approximately normal, with most participants clustering around 20:00 (**Figure 4A**). A nonlinear association with age was observed, with younger individuals displayed later chronotypes compared to older individuals (**Figure 4B**), consistent with previous findings using melatonin measurements of circadian phase (Kennaway, 2023). In addition, males exhibited slightly later chronotypes than females (**Figure 4C**). This sex difference was further supported by the model that additionally included age (**Figure 4C**, inset plot), suggesting an average difference of approximately 6 minutes.

**Figure 4.**
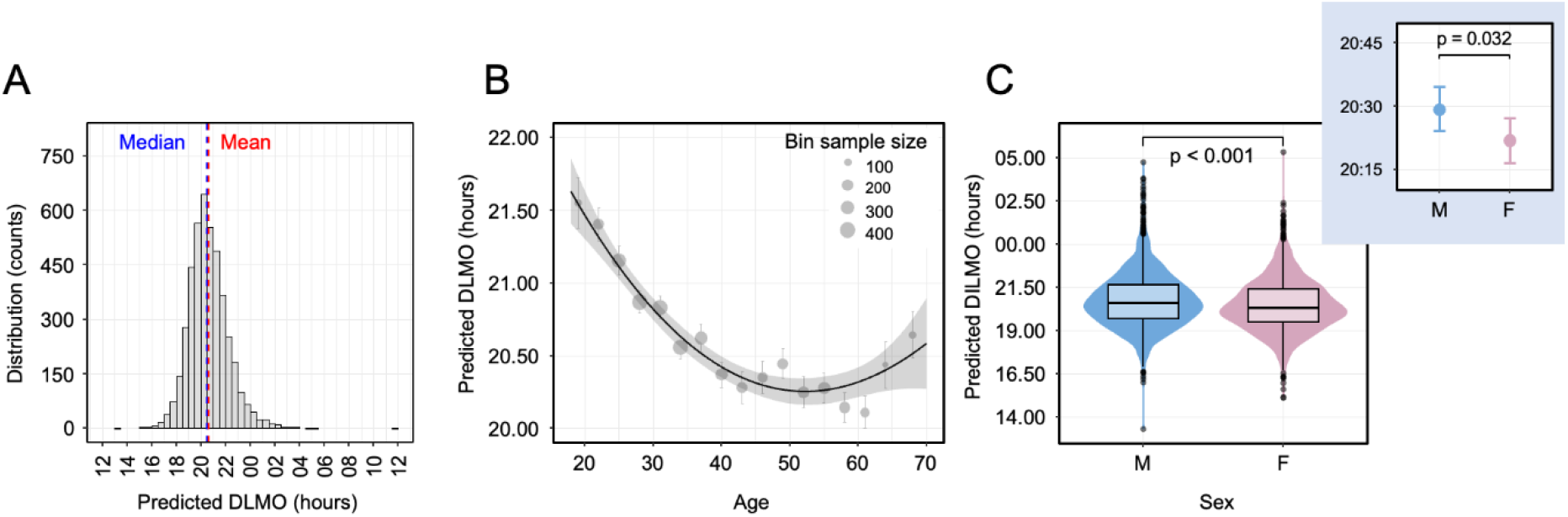
Epidemiology of human chronotype. (**A**) Distribution of chronotype in the entire sample (n = 4,351). Chronotype predicted from hair follicles is associated with age (**B**) and sex (**C**) (n = 3,949). In (A), the red dashed line represents the mean, and the blue dashed line represents the median. In (B), the line represents the fit from model A, with age as a spline term. The shaded area shows the 95% confidence interval of the fit. Dots and whiskers represent chronotype (raw data) mean ± standard error by age category. In (C), the main panel show sex differences, where p < 0.001 according to t-test. The insert depicts estimated marginal means and 95% confidence intervals from model A.

To investigate whether hair-based chronotype assessments reflect lifestyle-related differences, we conducted additional analyses based on the post hoc hypothesis that the slightly later chronotypes observed in participants over 60 years old could be attributed to a higher proportion of non-working individuals in that age group. As expected, participants who were occupationally active had an earlier chronotype than those not working (**Figure 5A**). Furthermore, participants who collected hair samples during the week (Wednesday/Thursday/ Friday) had an earlier chronotype than those sampled on Sunday (**Figure 5B**), suggesting a shift in phase of entrainment between weekdays and weekends. These findings were further supported by models adjusted for age and sex, which estimated that working individuals had a chronotype approximately 34 minutes earlier than non-working individuals, while weekday samples showed a chronotype about 12 minutes earlier than those collected on Sundays (**Figure 5A-B**, insets). Finally, the number of self-reported workdays per week was negatively correlated with chronotype (**Figure 5C**), with an earlier chronotype observed among individuals working more days per week. Taken together, these findings suggest that the human circadian system adapts to societal schedules, as reflected in occupational factors, thereby demonstrating its inherent plasticity.

**Figure 5.**
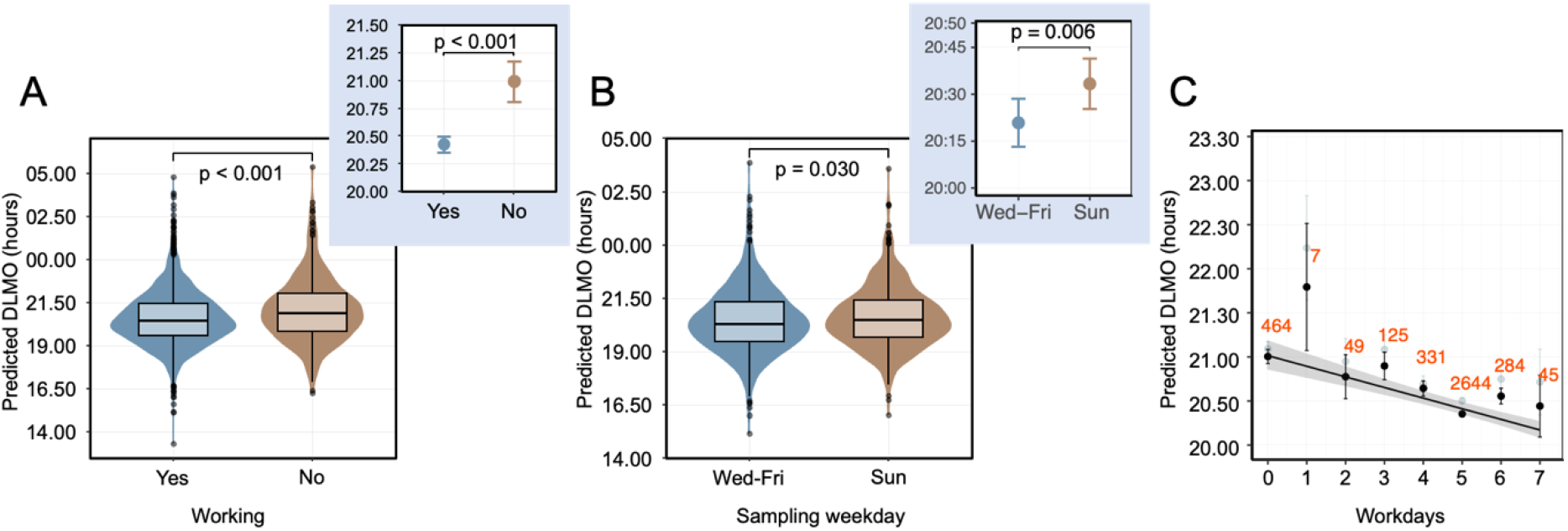
Predicted chronotype and work schedules. (**A**) Predicted DLMO (chronotype) is earlier among individuals who report being occupationally active compared to those who are not (n = 3,949, t-test). The inset plot shows estimated marginal means and 95% confidence intervals from model B, adjusted for age and sex. (**B**) Chronotype is also earlier when hair samples were collected on a weekday (Wednesday, Thursday, or Friday) compared to Sunday (n = 1,450, t-test). The inset plot displays estimated marginal means and 95% confidence intervals from model C, adjusted for age and sex. (**C**) A higher number of self-reported workdays per week is associated with earlier chronotype (n = 3,949). The line represents the fit from linear model D, adjusted for age and sex. Dots and whiskers show mean chronotype ± standard error, with raw data in gray and adjusted values in black. Sample sizes for each category are indicated in orange.

## Discussion

We developed and validated HairTime, a simple and accurate test for determining human chronotype, requiring only a few hair roots collected at a single time point during the day. Methods like this, which can be scaled and integrated into clinical routines, are essential for advancing our understanding of circadian rhythms and their role in health and disease. Accurate assessments of circadian phase of entrainment, or chronotype, are especially valuable for personalizing chronotherapy—helping determine the optimal timing for treatments or medications based on an individual’s internal time. The current gold standard for assessing chronotype, DLMO, is cumbersome, requiring multiple blood or saliva samples collected in a controlled environment prior to bedtime. Previously, we developed the BodyTime assay, which estimates chronotype using a single blood sample, but its collection and processing limitations restricted its use to smaller, laboratory-based studies. In contrast, HairTime is minimally invasive and provides similarly accurate predictions in both training and validation sets, with a median absolute deviation from saliva DLMO of less than 1 hour. Over 85% of the samples showed deviations of less than 2 hours. Our chronotype estimate is based on the average of predictions from three models—ZeitZeiger, LASSO, and PLS—and we demonstrate that the standard deviation of the model predictions can serve as an approximation of accuracy. Higher standard deviations between models correspond to poorer prediction accuracy. The machine learning models used selected molecular transcript biomarkers, many of which are genes associated with the core circadian oscillator (e.g., PERs, CRYs, ARNTL, NR1Ds). These genes have also been identified as biomarkers in other chronotype tests, where they were detected in biological samples like blood. However, we also observed genes specific to hair follicle cells, suggesting that a combination of widely expressed clock genes and tissue-specific clock output genes provides the optimal basis for accurate chronotype prediction

Recent studies have explored various methods to estimate the phase of entrainment using a single or few biological samples, such as gene expression and metabolites in blood, or gene expression in skin (Braun et al., 2018; del Olmo et al., 2022; Laing et al., 2017; Woelders et al., 2023; Wu et al., 2018). However, our study is the first to use hair follicles collected at a single timepoint as a source for chronotype estimation. Hair follicles offer several advantages as a biological material: they are more accessible, can be collected with minimal or no invasiveness, and are more stable than other samples like urine or blood. Moreover, hair samples are easier to store and transport, and their preparation is less expensive. Participants can collect hair samples in a stabilizing solution and easily mail them back at room temperature, making the process even more convenient for large-scale data collection. Our analysis, based on over 4,000 samples from the BodyClock dataset, demonstrates HairTime’s feasibility and scalability for larger-scale studies.

Since the prediction accuracy of HairTime varied between subjects, we hypothesized that the amplitude of circadian transcript rhythms might partly explain this variability. Specifically, we assumed that phase predictions are likely to be less accurate when the amplitude is lower. To explore this, we developed an amplitude score based on the rhythms of five circadian genes: CIART, TEF, NR1D1, NR1D2, and PER3. These genes were chosen due to their high amplitude and strong cross-correlation in the HairTime training sample, with the latter three genes previously identified as key indicators of circadian properties in hair follicles (Akashi et al., 2010). Interestingly, individuals with lower amplitude scores showed poorer agreement between the chronotype predictions of the three models in the training set. This finding supports our hypothesis that the accuracy of chronotype prediction is influenced by the strength of circadian rhythms in the hair follicle cells. To predict this amplitude score from a single timepoint, we identified six genes with minimal variation in expression throughout the day and trained a model to estimate amplitude based on their absolute expression levels in the morning. As expected, in the validation study, the predicted amplitude score correlated with the accuracy of chronotype predictions.

The concept of circadian amplitude is not clearly defined, and there is no consensus on a gold standard for its assessment. Reduced amplitudes in rest-activity and skin temperature rhythms have been associated with aging and various diseases (Brooks et al., 2023; Feng et al., 2023; Musiek et al., 2018). While it is commonly assumed that this decline reflects changes in the endogenous circadian system, it remains unclear whether such reductions are governed by the circadian system itself or are influenced by external factors (Dijk & Duffy, 2020). Nonetheless, the amplitudes of melatonin, cortisol, and core body temperature respond similarly to interventions, suggesting they may reflect the amplitude of suprachiasmatic nucleus (SCN) oscillations (Dijk et al., 2012). It is still unknown whether amplitude information derived from hair follicle cells can serve as a marker of the central clock’s amplitude or if it primarily reflects that of local clocks. Given the multi-oscillator nature of the human circadian system, the amplitudes and phases in peripheral tissues may, under certain conditions, reflect either the rhythmicity imposed by the central clock or that of local clocks—an area that warrants further investigation. In any case, our amplitude prediction model based on a single hair sample could be a valuable tool for exploring these questions. It may also serve as an effective means of tracking the success of interventions aimed at enhancing the amplitude of the circadian system, e.g. through light therapy, melatonin, or time-restricted eating (Kramer et al., 2022).

While our phase models were trained to predict DLMO, and thus the phase of the SCN, it remains unclear whether their accuracy is maintained when sleep patterns are displaced (e.g., in shift work or jetlag). For example, predictions of DLMO from activity and light exposure tend to be less accurate when individuals are misaligned (Brown et al., 2021). Mistimed sleep has been shown to affect the rhythmicity of circadian transcripts in whole blood samples (Archer et al., 2014), suggesting that expression profiles in peripheral tissues are not only influenced by the central circadian clock but also by external cues, such as temperature, nutrition, and those related to the sleep-wake cycles. Previous research showed that hair follicles phase correlate with the phase of behavior (i.e., sleep-wake, eating times). However, in an experiment where sleep-wake and eating times were advanced by 4 hours over 3 weeks, along with high-intensity light exposure to adjust circadian rhythms, the phase shifts in hair follicle transcript rhythms were more closely aligned with those observed in salivary melatonin and cortisol than with those in sleep behavior (Akashi et al., 2010). Additionally, the phase shifts in hair follicle rhythms were less drastic compared to activity rhythms in rotating shift workers. These findings suggest that hair follicle phase is not highly susceptible to mistimed sleep and that even in challenging situations like sleep displacement, the HairTime assay may still provide relatively accurate estimates of circadian phase, making it a valuable tool for assessing phase in conditions where the sleep-wake cycle is disrupted.

In the large BodyClock dataset, the distribution of chronotype estimated using the HairTime assay mirrored the patterns observed in DLMO, including its well-documented age dependency (Kennaway, 2023). However, sex differences remain unclear and inconsistent across studies. For example, while they may not be evident in DLMO and MEQ (Kennaway, 2023), men were slightly later when chronotype was assessed using the MCTQ (Roenneberg et al., 2007). Recent studies have suggested that chronotype differences between men and women may be smaller than previously thought (Randler & Engelke, 2019). Consistent with these findings, we observed that chronotype predicted from hair samples was slightly later in men. This suggests that HairTime can reliably capture age- and sex-related chronotype variations, similar to other established measures such as DLMO, but also offers the potential for large-scale epidemiological studies of circadian rhythms.

Interestingly, our exploratory analyses with the BodyClock dataset suggest some degree of influence of lifestyle and environmental factors on hair-based chronotype predictions. For example, samples collected on weekends revealed generally later chronotypes, while subjects who reported working more days had an earlier chronotype. This highlights the plasticity of chronotype, which is shaped not only by the endogenous circadian system but also by behavioral and environmental factors. A critical point is that chronotype, often mistakenly viewed as a purely genetic trait or behavioral preference, should instead be understood as a dynamic state—resulting from the complex interaction between the genetically determined circadian system and external environmental cues. These factors together establish the phase of entrainment, or chronotype. Thus, it is crucial to recognize that chronotype is not a fixed characteristic dictated solely by genetic makeup; rather, it is a dynamic state that can vary in response to external influences like light or food (zeitgeber exposure), which can be shaped by behavioral factors such as work schedules, sleep patterns, mealtimes, and time spent outdoors or in front of screens.

A key question that remains is how much of this observed plasticity reflects changes in the internal circadian clock, as opposed to behavior deviating from the internal clock phase and influencing local gene expression. Additionally, a related question is which of the two explains the negative health outcomes associated with social jetlag and sleep timing irregularity: an internal clock that is constantly shifting or behaviors that constantly shift and are misaligned with the clock? Recent research has shown that DLMO is less stable than previously believed, with shifts in the phase of entrainment observed between workdays and work-free days, especially in late chronotypes (Zerbini et al., 2021). This finding raises the possibility that the phase of the central clock may be more plastic than previously thought, or that DLMO, as an overt rhythmic output, may be less robust than the central clock itself. These considerations highlight the need to explore the mechanisms underlying circadian plasticity, especially as they relate to health outcomes. Evidence suggests that interventions should aim to stabilize both behavioral and internal rhythms to mitigate the health risks associated with social jetlag and sleep timing irregularities (Chellappa et al., 2021; De Sá Couto-Pereira et al., 2024; Sletten et al., 2023). The difference in internal phase estimated by HairTime between week- and weekend days was around 12 minutes, while Zerbini et al. observed a larger difference in DLMO of nearly 30 minutes between work- and work-free days in their study. This discrepancy may be due to differences in study design, as Zerbini et al. used a within-subjects approach in a smaller sample. Nevertheless, HairTime offers a powerful tool for examining chronotype stability across time, enabling future studies to track its changes and correlations with sleep patterns, providing new insights into circadian plasticity and disruptions.

Our results should be interpreted in light of the study’s strengths and limitations: (i) HairTime was developed using a training set and validated on a separate validation set, demonstrating good accuracy compared to the gold standard, DLMO. The assay relies on fewer than 20 transcript biomarkers and combines three models to predict chronotype, with the agreement between models serving as an indicator of accuracy. However, it is important to note that the models were trained on a relatively small cohort, which may limit their generalizability. Additionally, the association between lower amplitudes and higher prediction error suggests that future studies should evaluate HairTime’s accuracy in individuals with mistimed sleep in controlled settings (e.g., manipulated rest-activity rhythms). Further testing in men and women with diverse genetic backgrounds, as well as in relevant patient groups, is warranted. (ii) Our analyses using the BodyClock dataset strongly support the applicability of HairTime in field studies, given the large sample size and the inclusion of various age groups and work characteristics. Although data were collected via convenience sampling and analyses were exploratory, we observed demographic patterns in hair-based chronotype that align with those reported in the literature. Unfortunately, we lacked information on work start times, which could have provided a more complete understanding of the relationship between hair-based internal time and work schedules. Our comparisons of weekdays vs. weekends were cross-sectional; despite controlling for age and sex and including only participants with 5 workdays per week, future studies using a within-subject design should further validate these findings.

In conclusion, HairTime is a pioneering, non-invasive assay that estimates chronotype from hair samples collected at one time point during the day, making it an ideal tool for large-scale studies, particularly in field settings. The ability for participants or patients to self-collect samples, combined with their stability and ease of transport, ensures broad applicability. Our analyses in the large BodyClock cohort reveal significant insights into the plasticity of chronotype, demonstrating how chronotype, as measured by HairTime, adapts to different environmental and behavioral factors such as work schedules. These findings suggest that chronotype is not a fixed trait but a dynamic state that can shift based on external influences. The ability to measure this plasticity is crucial for circadian medicine, as it enables personalized treatment approaches and interventions. HairTime can be easily collected longitudinally to track individual chronotype fluctuations over time allowing for more precise tailoring of therapies. Such longitudinal assessments can advance our understanding of how circadian rhythms adapt to different conditions and how they can be optimized for better health outcomes.

## Materials and Methods

### Ethical approval and informed consent

The studies in this paper were approved by the Ethics Committee of Charité – Universitätsmedizin Berlin (EA1/330/19 and EA4/200/19). All participants provided written informed consent prior to participation, acknowledging their voluntary involvement and understanding of the study procedures.

### Pilot study

Three healthy volunteers (2 males, 1 female) collected hair follicle samples every 4 hours over a period of 36 hours (10 samples per donor). RNA was isolated from the samples, which were then sent for RNA sequencing (Novogene). Rhythmic gene expression was identified using MetaCycle (Wu et al., 2016). Genes were only included if at least 50% of the samples had an expression value of at least 1 for each of the three subjects resulting in 14,049 genes analyzed. Based on rhythmicity and amplitude, 90 candidate gene transcripts and 6 housekeeping gene transcripts were selected as potential time markers for the *HairTime* study. For details on sequencing quality, read numbers, and selection criteria, please refer to **Supplementary Table 1**.

### HairTime study

This study aimed to identify transcript biomarkers in human hair follicles for predicting chronotype. Thirteen healthy subjects were recruited (**Supplementary Table 2**) and trained on proper hair follicle sampling techniques. Participants were instructed to maintain a regular sleep-wake schedule for 7 days prior to sample collection, with compliance monitored using activity watches (Motion Watch 8, CamNtech) and sleep diaries. On day 8, participants collected 10 hair follicles every 3 hours over a 33-hour period (12 samples). Six hours before their habitual bedtime on day 9, participants visited the lab for melatonin saliva sampling at 30-minute intervals. Dim light melatonin onset (DLMO) was determined as detailed below. RNA was isolated from hair samples, and gene expression was quantified as detailed below. The expression profiles of 96 genes (**Supplementary Table 3**) were analyzed using the nCounter platform from NanoString.

### HairVali study

The aim of this study was to validate the transcript biomarkers identified in HairTime using an independent cohort. Thirty-five healthy volunteers (**Supplementary Table 2**) were instructed to maintain a regular sleep-wake schedule for 7 days prior to sampling. On the morning of day 8, participants visited the lab to donate a hair follicle sample. In the evening of the same day, they returned to the lab 6 hours before their usual bedtime for saliva sample collection to determine DLMO.

### Hair Follicle Sampling

Participants were trained in hair follicle collection and asked to practice prior to starting. Briefly, they were instructed to grasp individual hairs or small strands with their fingers or tweezers and pull sharply in the direction of growth. After collection, participants visually confirmed that follicles had been extracted and placed them in collection tubes containing 1.5 ml RNAlater, prepared for each time point. Once the time series was completed, samples were transported to the lab on the same day and stored at 4 °C until further processing.

### RNA isolation from hair follicles

was performed using a TRIzol-based solid phase extraction method. Briefly, RNAlater solution was removed, and 800 µl of TRIzol was added directly to the tube containing the hair follicle sample. The sample was vortexed briefly and incubated for 5 minutes at room temperature. RNA was isolated following the protocol of the Direct-Zol RNA Micro Kit (Zymo Research). The RNA was then eluted in 10 µl of RNase-free water, stored on ice until quantification with the Qubit RNA Broad Range Assay Kit (Invitrogen), and stored at -80 °C for long-term preservation.

### DLMO assessment

Subjects entered a dimly lit room (< 5 lx) 6 hours before their habitual bedtime. Saliva samples were collected every 30 minutes (total of 12 samples per subject) using Salivettes (Sarstedt AG & Co.). The first sample was taken after 30 minutes under dim-light conditions, with the last sample collected at habitual bedtime. Eating and drinking were prohibited for 20 minutes prior to sampling. Upon collection, each sample was stored at –20 °C. Melatonin levels were measured using the Direct Saliva Melatonin Kit (BÜHLMANN). Circadian phase was determined by calculating the dim-light melatonin onset (DLMO) using the threshold method in Hockey-stick software (Danilenko et al., 2014), with a melatonin onset threshold of 3 pg/ml.

### Quantification of gene expression

A custom 96-plex NanoString panel was designed, consisting of 90 candidate time-telling genes and 6 housekeeping genes (**Supplementary Table 3**). The probes included a 3′-biotinylated capture probe and a 5′-fluorescence-barcoded reporter probe for each target gene. This panel was used in the HairTime and SleepShiftHairVali studies. Probe hybridization was performed with 400 ng of RNA from hair follicles, following the manufacturer’s instructions. Raw expression data were obtained using the NanoString nCounter Digital Analyzer (NanoString Technologies). Counts were normalized to the housekeeping gene *CTLC* and log2-transformed for downstream analyses. A reduced panel of 24 time-telling, amplitude, and housekeeping genes was used to analyze samples from the BodyClock cohort (**Supplementary Table 5**).

### Prediction of DLMO from hair follicle gene expression panel

In a recent study, we applied three machine learning methods—LASSO, PLS, and ZeitZeiger (Franklin, 2005; Hughey et al., 2016; Wittenbrink et al., 2018))—to predict DLMO from transcriptome data of peripheral blood monocytes. In this study, we applied the same approach to hair follicle samples. Briefly, a training dataset was generated using gene expression data from the NanoString nCounter platform, collected during the HairTime study. These 13 time-series were time-stamped according to the individual time relative to DLMO. LASSO, PLS, and ZeitZeiger were then used in the R statistical environment (R Core Team, 2024) to identify predictors from the time-series data.

### BodyClock dataset – preprocessing

The HairTime biomarkers described in this study have been patent-pending (EP20210217005) and licensed to BodyClock Technologies, a spin-off from Charité Universitätsmedizin Berlin. BodyClock offers the test in a self-sampling kit as a lifestyle product, which includes detailed instructions for hair follicle sampling. After sampling, customers are instructed to register their sample online and send it by mail to the Laboratory of Chronobiology at Charité. During registration, customers are asked to complete a brief online questionnaire about their sleep habits and occupational status. BodyClock shared the anonymized and aggregated data for research purposes, ensuring their data is processed in accordance with applicable data protection laws. Between December 2021 and June 2024, this process resulted in the collection of the BodyClock dataset, the first analysis of which is presented in this paper.

Data from questionnaires and assays were merged and preprocessed using R (v. 4.2.2), *tidyverse* (v. 1.1.2, (Wickham et al., 2023)) and *janitor* (Firke, 2024). During the merging process, extra entries were removed using criteria based on the following: quality control annotations, proximity of questionnaire submission and hair sampling dates, and questionnaire data plausibility. This resulted in the removal of 108 extra assay entries and 311 questionnaire extra entries (with same id). Sensitivity analyses excluding any entry that had a duplicate showed similar results to those reported here (data not shown).

Entries where the assay failed quality control were excluded when assessing the distribution of chronotype (n= 180). For analyzing the association of chronotype with age, sex and work, we included only subjects with reliable self-reported data on age and sex (excluding n = 296) and work (excluding n = 11). Data from participants younger than 18 or older than 70 were also excluded (n = 95) due to underrepresentation, which could yield unreliable results.

The self-reported date of hair sampling was used to investigate chronotype differences by weekday. We included only participants who collected their hair samples on Wednesday, Thursday, Friday, or Sunday (excluding n = 1,773). Additionally, we restricted the analysis to those who reported working 5 days a week (excluding n = 726). Our assumption was that most subjects working 5 days/week work Monday through Friday and that transition days may carry residual effects of the weekend or the week.

Sample sizes for each analysis are provided in the results section and figure legends. **Supplementary Figure 1** shows the dataset sample sizes and exclusion counts.

### BodyClock dataset – data analysis

Given the nearly normal distribution of chronotype, we used t-tests to examine the association between chronotype and categorical variables (sex, workstatus and weekdays vs. weekend) and Pearson’s correlation to examine chronotype association with age and workday. To assess their independent effects, we performed a multivariable regression (model A) adjusting for both age and sex. We then analyzed the influence of work status (yes vs. no) and weekday (weekday vs. weekend) by first comparing chronotype between groups using t-tests, followed by multivariable regressions (models B and C) adjusted for age and sex. Finally, we assessed whether the number of workdays was associated with chronotype in a multivariable regression adjusted for age and sex (model D). All t-tests, Pearson correlations, and linear models were conducted using Base R (v. 4.2.2). Since our analyses are exploratory, p-values should be interpreted accordingly. To extract regression results, we used the package *jtools* (v. 2.3.0, (Long, 2024)) and *ggeffects* (v. 1.7.1, (Lüdecke, 2018)) to derive marginal means (predicted values) for visualization. We used *ggplot2* (v. 3.5.1, (Wickham, 2009)), *ggpubr* (v. 0.6.0, (Kassambara, 2023)) for plotting. For linear independent variables of interest in each model (i.e., age and workdays), we used Bayesian information criterion (BIC) to compare two functional forms: linear and spline transformation using *splines::bs* (v. 4.4.2, (Bates & Venables, 2021)). We opted to use BIC, a Bayesian approach, as it provides a balance between model fit and model complexity, penalizing models with more parameters and thus helping to prevent overfitting. The model with the lower BIC value was selected as the best fit, and the selected model is reported in each case. The R code for the plots, tests and model results, and assumptions checks (linearity, normality of residuals with Q-Q plots, and homoscedasticity with residual vs. fitted plots) is available at [github.com/Chronomedicine/HairTime.git]. All analyses should be considered exploratory, and the results should be interpreted accordingly.

## Author contribution

Defined according to contributor role taxonomy (*CRediT*, n.d.). B.M.: Conceptualization (equal); data curation (equal); formal analysis (supporting); investigation (equal); methodology (equal); supervision (equal); writing – original draft (supporting); writing – review&editing (equal). S.Ö.: Investigation (equal); methodology (equal); writing – review&editing (supporting). L.K.P.: Conceptualization (equal); data curation (lead); formal analysis (lead); investigation (equal); writing – original draft (lead); writing - review&editing (equal). A.R.: data curation (equal); formal analysis (supporting); writing - original draft (supporting), writing – review&editing (supporting). A.A.: Investigation (supporting); methodology (supporting); writing – review&editing (supporting). J.Z.: Investigation (supporting); writing – review&editing (supporting). D.K.: Project administration (equal); supervision (equal); writing - review&editing (supporting). A.K.: Conceptualization (lead); data curation (lead); formal analysis (lead); methodology (equal); funding acquisition (lead); project administration (lead); supervision (lead); writing – original draft (lead); writing – review&editing (equal).

## Acknowledgements

We thank Dr. Theresa Keller for reviewing our code and text on methods/results of BodyClock analyses.

