## Supplementary figures and images for "Hair test reveals plasticity of human chronotype"

### Supplementary Figure 1

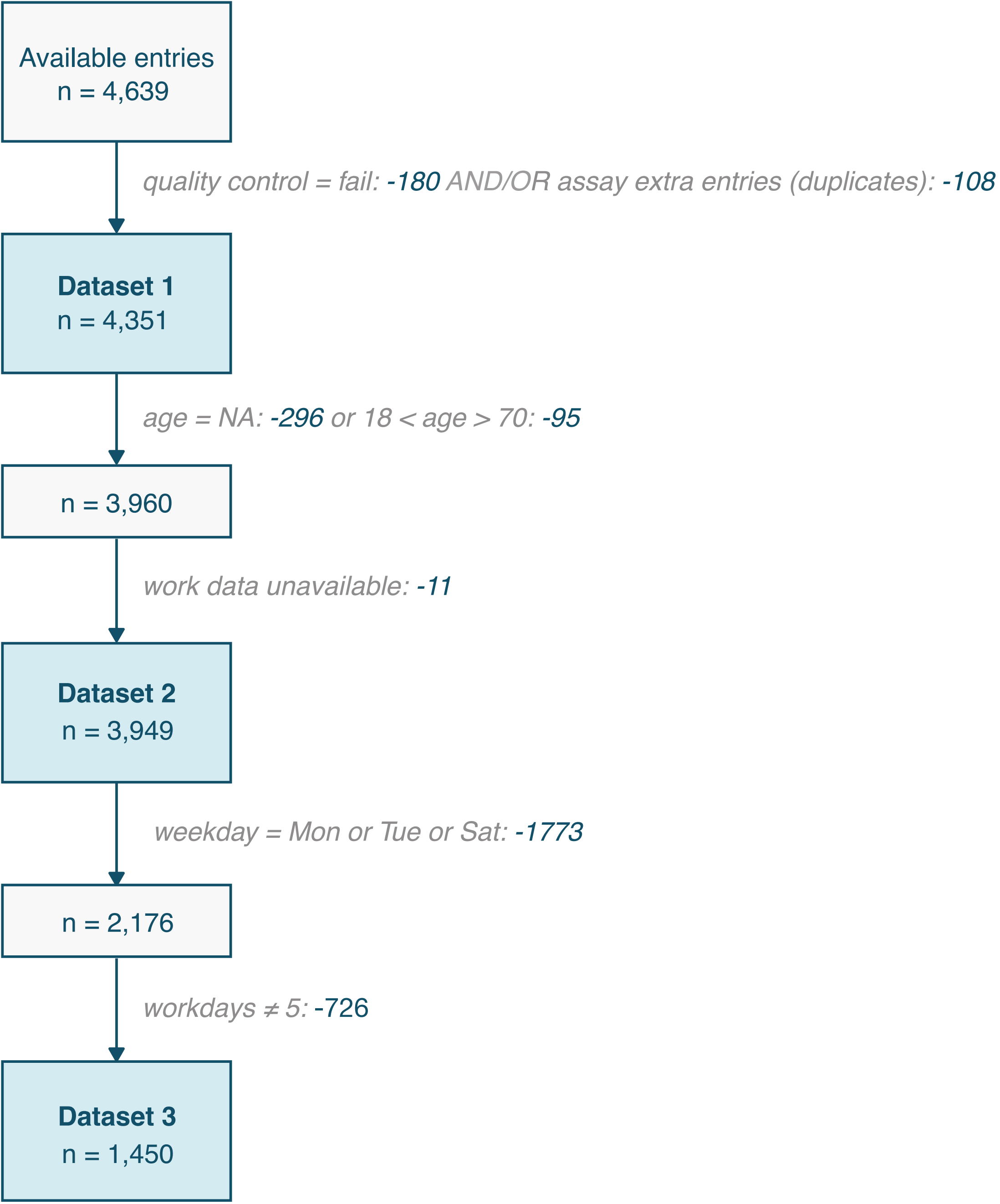
