## Supplementary Table 2 for "Hair test reveals plasticity of human chronotype"

**Supplementary Table 2: HairTime and HairVali studies – Participants’ age, scores of PSQI and MCTQ (MSF-sc)^a^ as well as habitual bedtimes (BT) and DLMO times based on the threshold method^b^.**

|  | **Age** | **PSQI** | **MCTQ**  **(MSF-sc)** | **BT** | **DLMO**  **(threshold)** |
| --- | --- | --- | --- | --- | --- |
| **HairTime**  **All (n=13)** | 31.1 (7.7)  22-52 | 4.6 (2.0)  1-8 | 4.8 (1.0)  2.5-6.3 | 00:03 (0:58)  22:05-1:30 | 21:45 (0:59)  20:21-23:31 |
| **HairTime**  **Females (n=6)** | 27.2 (3.4)  22-31 | 4.7 (2.2)  1-7 | 4.8 (0.8)  3.8-6.0 | 00:12 (0:44)  23:10-1:30 | 21:34 (1:02)  20:21-23:01 |
| **HairTime**  **Males (n=7)** | 34.4 (8.9)  25-52 | 4.6 (2.0)  2-8 | 4.8 (1.2)  2.5-6.3 | 23:55 (1:10)  22:05-1:30 | 21:54 (1:00)  20:35-23:31 |
| **HairVali**  **All (n=35)** | 37.3(12.2)  20-64 | 4.6 (1.3)  1-10 | 4.4 (1.4)  0.9-7.0 | 23:31 (01:00)  21:30-02:15 | 20:40 (1:18)  16:54-23:11 |
| **HairVali**  **Females (n=21)** | 38.5 (13.2)  20-64 | 4.9 (2.2)  1-10 | 4.1 (1.4)  0.9-6.1 | 23:23 (00:54)  21:52.00:53 | 20:18 (1:13)  16:54-22:31 |
| **HairVali**  **Males (n=14)** | 35.4 (10.7)  22-62 | 4.2 (1.3)  2-6 | 4.8 (1.4)  1.6-7.0 | 23:44 (01:07)  21:30-02:15 | 21:13 (1:19)  18:42-23:11 |

^a^ Local time of mid-sleep on free days corrected for sleep debt accumulated over the workweek.

^b^ Given are mean scores, SD (in brackets) and ranges. BT and DLMO are given in clock time (hh:mm).
