## Supplementary Table 5 for "Hair test reveals plasticity of human chronotype"

**Supplementary Table 5.** Gene sets used to determine expression profiles of 96- and 24-time telling genes including housekeeping genes on the nCounter platform from NanoString (Bruker).

|  | HairTime 96plex | HairTime 24plex |
| --- | --- | --- |
| Time Telling genes | ARNTL AC011476.3 AC016727.1 AC048341.2 AL391121.1 ALDH3A1 ANO7L1 ARSJ B3GNT2 BORCS6 C12orf66 C1orf123 CAMKMT CD34 CDC25B CDCA3 CDPF1 CIART CLEC11A CNN1 CRY1 CRY2 DBP DIO3OS DUBR FBLN7 GLI2 GNB1L GOT1 HERC3 HLF KLRG2 KRT15 LAYN LRAT LRRC37A3 MAP3K14 MINOS1 MOSPD2 NDUFAF4 NME7 NPAS2 NR1D1 NR1D2 NR1H3 P4HA1 PDE6B PDK1 PDSS1 PER1 PER2 PER3 PGF PIBF1* PIF1 PLEKHF1* POLR2I* POLR2J4 POLR3GL* PRR34-AS1* PSMG3-AS1* RAI14* RBM18* RORγ* RPL23AP7 RRAD SAP30 SCGB2A2 SERPINE2 SLC22A15 SLC6A6 SPRR2A SPRR2B SPRY2 STXBP4 TDRKH TEF TGM5 TLR5 TNS1 TRIM35 U2AF1 URB1-AS1 VWF ZBTB42 ZNF296 ZNF510 ZNF669 ZNF749 ZSCAN31 | ARNTL CIART CRY1 CRY2 DBP DUBR GLI2 KRT15 MOSPD2 NR1D1 NR1D2 PDK1 PER1 PER2 PER3 PGF SPRY2 TEF TGM5 TLR5 TRIM35 ZNF296 ZNF749 |
| Housekeeping genes | CLTC GAPDH HPRT1 PGK1* PPIA* PSMB2* | CLTC |

*Genes excluded from analysis due to manufacturing issues.
