## Supplementary Table 6 for "Hair test reveals plasticity of human chronotype"

**Supplementary Table 7:** BodyClock-derived datasets

|  | **Dataset 1** (N=4351) | **Dataset 2** (N=3949) | **Dataset 3** (N=1450) |
| --- | --- | --- | --- |
| **Age category** |  |  |  |
| <18 | 1 (0.0%) | 0 (0%) | 0 (0%) |
| 18-25 | 523 (12.0%) | 521 (13.2%) | 167 (11.5%) |
| 26-50 | 2383 (54.8%) | 2380 (60.3%) | 960 (66.2%) |
| 51-70 | 1054 (24.2%) | 1048 (26.5%) | 323 (22.3%) |
| >70 | 94 (2.2%) | 0 (0%) | 0 (0%) |
| Missing | 296 (6.8%) | 0 (0%) | 0 (0%) |
| **Sex** |  |  |  |
| M | 2119 (48.7%) | 2073 (52.5%) | 803 (55.4%) |
| F | 1947 (44.7%) | 1876 (47.5%) | 647 (44.6%) |
| Missing | 285 (6.6%) | 0 (0%) | 0 (0%) |
| **Occupation** |  |  |  |
| No | 550 (12.6%) | 464 (11.7%) | 0 (0%) |
| Yes | 3504 (80.5%) | 3485 (88.3%) | 1450 (100%) |
| Missing | 297 (6.8%) | 0 (0%) | 0 (0%) |
| **Number of workdays** |  |  |  |
| Mean (SD) | 4.23 (1.80) | 4.32 (1.71) | 5.00 (0) |
| Median [Min, Max] | 5.00 [0, 7.00] | 5.00 [0, 7.00] | 5.00 [5.00, 5.00] |
| Missing | 297 (6.8%) | 0 (0%) | 0 (0%) |
| **Country of sampling** |  |  |  |
| Austria | 194 (4.5%) | 191 (4.8%) | 54 (3.7%) |
| Belgium | 3 (0.1%) | 3 (0.1%) | 1 (0.1%) |
| Denmark | 11 (0.3%) | 11 (0.3%) | 8 (0.6%) |
| Germany | 2879 (66.2%) | 2821 (71.4%) | 1051 (72.5%) |
| Italy | 4 (0.1%) | 4 (0.1%) | 0 (0%) |
| Liechtenstein | 3 (0.1%) | 3 (0.1%) | 0 (0%) |
| Luxembourg | 7 (0.2%) | 7 (0.2%) | 3 (0.2%) |
| Netherlands | 4 (0.1%) | 4 (0.1%) | 2 (0.1%) |
| Other country | 3 (0.1%) | 3 (0.1%) | 3 (0.2%) |
| Spain | 2 (0.0%) | 2 (0.1%) | 1 (0.1%) |
| Sweden | 2 (0.0%) | 2 (0.1%) | 0 (0%) |
| Switzerland | 104 (2.4%) | 99 (2.5%) | 35 (2.4%) |
| Missing | 1135 (26.1%) | 799 (20.2%) | 292 (20.1%) |
| **Year** |  |  |  |
| 2021 | 1 (0.0%) | 1 (0.0%) | 1 (0.1%) |
| 2022 | 1638 (37.6%) | 1467 (37.1%) | 528 (36.4%) |
| 2023 | 1534 (35.3%) | 1367 (34.6%) | 493 (34.0%) |
| 2024 | 1178 (27.1%) | 1114 (28.2%) | 428 (29.5%) |
| **Month** |  |  |  |
| Jan | 859 (19.7%) | 804 (20.4%) | 324 (22.3%) |
| Feb | 457 (10.5%) | 426 (10.8%) | 144 (9.9%) |
| Mar | 516 (11.9%) | 467 (11.8%) | 180 (12.4%) |
| Apr | 481 (11.1%) | 436 (11.0%) | 162 (11.2%) |
| May | 421 (9.7%) | 373 (9.4%) | 132 (9.1%) |
| Jun | 265 (6.1%) | 233 (5.9%) | 87 (6.0%) |
| Jul | 229 (5.3%) | 199 (5.0%) | 62 (4.3%) |
| Aug | 150 (3.4%) | 132 (3.3%) | 54 (3.7%) |
| Sep | 164 (3.8%) | 142 (3.6%) | 54 (3.7%) |
| Oct | 208 (4.8%) | 176 (4.5%) | 58 (4.0%) |
| Nov | 342 (7.9%) | 316 (8.0%) | 106 (7.3%) |
| Dec | 259 (6.0%) | 245 (6.2%) | 87 (6.0%) |
| **Weekday of hair sampling** |  |  |  |
| Friday | 454 (10.4%) | 420 (10.6%) | 253 (17.4%) |
| Monday | 704 (16.2%) | 645 (16.3%) | 0 (0%) |
| Saturday | 662 (15.2%) | 603 (15.3%) | 0 (0%) |
| Sunday | 987 (22.7%) | 905 (22.9%) | 662 (45.7%) |
| Thursday | 476 (10.9%) | 420 (10.6%) | 266 (18.3%) |
| Tuesday | 584 (13.4%) | 525 (13.3%) | 0 (0%) |
| Wednesday | 484 (11.1%) | 431 (10.9%) | 269 (18.6%) |
| **Hair sampling time** |  |  |  |
| Afternoon | 224 (5.1%) | 206 (5.2%) | 67 (4.6%) |
| Morning | 4127 (94.9%) | 3743 (94.8%) | 1383 (95.4%) |
